## Supplementary figures and images for "Development of diagnostic SNP markers for quality assurance and control in sweetpotato [*Ipomoea batatas* (L.) Lam.] breeding programs"

### Online Resource 3

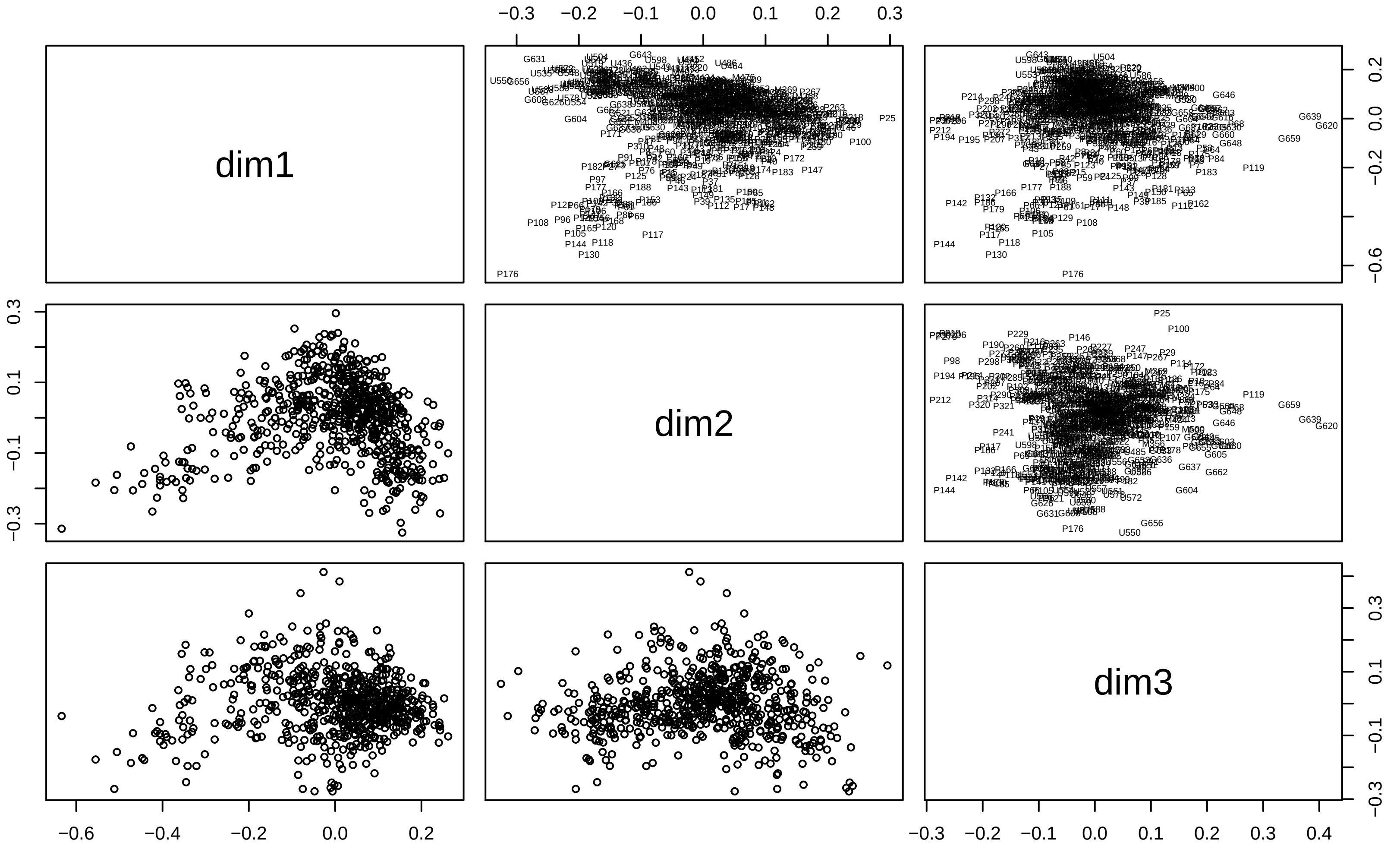

### Online Resource 4

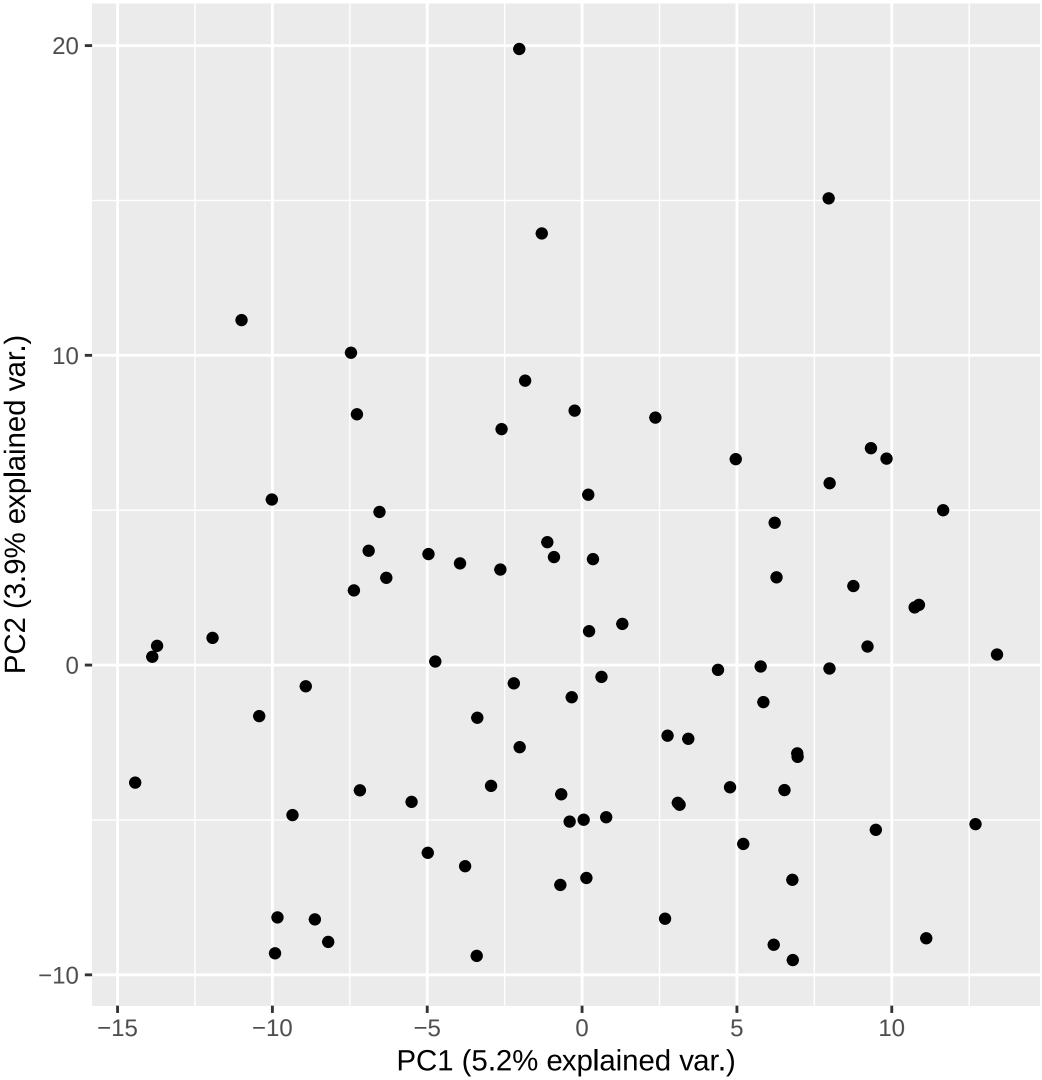
